# Type IV pili is a critical virulence factor in clinical isolates of *Paenibacillus thiaminolyticus*

**DOI:** 10.1101/2022.01.24.477451

**Authors:** Christine Hehnly, Aiqin Shi, Paddy Ssentongo, Lijun Zhang, Albert Isaacs, Sarah U. Morton, Nicholas Streck, Petra Erdmann-Gilmore, Igor Tolstoy, R. Reid Townsend, David D. Limbrick, Joseph N. Paulson, Jessica E. Ericson, Michael Y. Galperin, Steven J. Schiff, James R. Broach

## Abstract

Hydrocephalus, the leading indication for childhood neurosurgery worldwide, is particularly prevalent in low-and-middle-income countries (LMICs). Hydrocephalus preceded by an infection, or postinfectious hydrocephalus (PIH), accounts for up to 60% of hydrocephalus in LMICs. Since many children with hydrocephalus suffer poor long-term outcomes despite surgical intervention, prevention of hydrocephalus remains paramount. Our previous studies implicated a novel bacterial pathogen, *Paenibacillus thiaminolyticus,* as a contributor to PIH in Uganda. Here we report the isolation of three *P. thiaminolyticus* strains, Mbale, Mbale2, and Mbale3, from patients with PIH and the demonstration that the three clinical isolates exhibit virulence in mice while *P. thiaminolyticus* type strain, B-4156, does not. We constructed complete genome assemblies of the clinical isolates as well as the reference strain and performed comparative genomics and proteomics analyses to identify potential virulence factors. One candidate virulence factor is a cluster of genes carried on a mobile genetic element that encodes a type IV pilus and is present in all three PIH patient strains but absent in the type strain. Proteomic and transcriptomic data confirmed the expression of this cluster of genes in the Mbale strain, while CRISPR-mediated deletion of the gene cluster substantially reduced the virulence of this strain. Our comparative proteogenomic analysis also identified various antibiotic resistance loci in the virulent strains. These results provide insight into the mechanism of virulence of *Paenibacillus thiaminolyticus* and suggest avenues for the diagnosis and treatment of this novel bacterial pathogen.

**Author Summary:** Postinfectious hydrocephalus (PIH), a devastating sequela of neonatal infection, is associated with increased childhood mortality and morbidity. *Paenibacillus thiaminolyticus* was recently identified as the dominant organism highly associated with PIH in an African cohort. Our whole-genome sequencing, RNA sequencing and proteomics of three clinical isolates and a type strain in combination with CRISPR editing has revealed the type IV pili (T4P), encoded in a mobile genetic element, as a critical virulence factor for *P. thiaminolyticus* infection. Given the widespread presence of T4P in pathogens, the presence of T4P operon could serve as an important diagnostic and therapeutic target in *P. thiaminolyticus* and related bacteria.

## Introduction

Hydrocephalus is one of the most common brain disorders in children globally [1, 2] and the most common indication for pediatric neurosurgery [2]. A serious infection such as neonatal sepsis often precedes hydrocephalus [3], and postinfectious hydrocephalus (PIH) accounts for up to 60% of the nearly 400,000 children who develop hydrocephalus each year, principally in low-and middle-income countries (LMICs) [1, 3, 4]. PIH remains the leading cause of neurological morbidity and mortality worldwide despite recent clinical efforts to optimize the treatment [5, 6].

Strategies to prevent PIH have been thwarted for two principal reasons. First, standard clinical evaluation often fails to identify the pathogen(s) responsible for the underlying infectious episodes that precede PIH [7], precluding optimized or targeted treatment of the underlying infections. Second, the underlying pathophysiologic changes that lead to hydrocephalus following infection remain unknown [8]. Unfortunately, even children who undergo surgical treatment for hydrocephalus early in life can suffer very poor long-term outcomes [6, 9]. Therefore, major advances in the health of these children will require preventing infection by targeting both the underlying pathogens and their routes of infection [10–12], as well as improving treating infections with more optimal antibiotics and adjunctive therapies that can reduce the likelihood of subsequently developing hydrocephalus.

We have identified and isolated a novel pathogenic bacterial strain, *Paenibacillus thiaminolyticus* Mbale, as a likely causative agent in a significant fraction of a cohort of PIH cases in Uganda [13]. We showed that *P. thiaminolyticus* Mbale is lethal to mice following peritoneal injection, whereas injection of the *P. thiaminolyticus* type strain, NRRL^T^ B-4156 (B-4156), does not cause lethality. *Paenibacillus* species have been isolated and studied from various sources, particularly in agricultural and industrial applications [14]. Although some species such as *P. alvei* and *P. larvae* are known to cause widespread disease in honeybees [15], until recently only anecdotal cases of human disease associated with *Paenibacillus* have been reported [16–21]. We recently confirmed our initial report implicating *P. thiaminolyticus* as a causative agent of PIH by identifying *P. thiaminolyticus* infection in the CSF of 41% of 205 infants with PIH. Moreover, we detected *P. thiaminolyticus* in CSF samples from a significant fraction of neonates with clinical sepsis (Morton et al., unpublished data). Finally, we have isolated two additional clinical *P. thiaminolyticus* strains from patients in the larger PIH cohort and whose properties are described in this report.

Pathogenic bacteria typically have specific proteins, or virulence factors (VFs), that aid in their ability to survive and propagate in their hosts. Given the differential virulence between the B-4156 strain and our clinical isolates, we hypothesized that clinical isolates from our patients carry VFs that are absent in the nonpathogenic B-4156 strain. To address this hypothesis, we used genomic analysis to compare the predicted proteome of several clinical isolates to the B-4156 type strain and confirmed gene expression with RNA sequencing and proteomic analysis (**Fig S1).** For functional confirmation, we developed a novel CRISPR-Cas9 gene deletion system and applied it to one of the clinical isolates to confirm that a type IV pilus (T4P) identified with comparative proteogenomics is a critical VF. These results provide insight into the mechanism of virulence of *P. thiaminolyticus* and suggest avenues for diagnosis and treatment of this novel bacterial pathogen.

## Materials and Methods

The *Paenibacillus thiaminolyticus* type strain NRRL B-4156^T^ was obtained from the United States Department of Agriculture Agricultural Research Service Culture Collection (NRRL, Peoria, IL).

### Clinical Microbiology

Isolation and genome assembly of the Mbale strain were previously described [13]. To recover additional clinical isolates, 1 ml aliquots of CSF from sixty-three PIH patients [13] were inoculated into BD BACTEC^TM^ Lytic Anaerobic Medium culture bottles supplemented with 1 ml of defibrinated horse blood (Thermo Scientific). Culture bottles were incubated in a BD BACTEC^TM^ FX instrument and monitored for bacterial growth for up to 14 days. Culture bottles that were positive for bacterial growth were subcultured in BD BBL^TM^ Chocolate II and CDC Anaerobe 5% Sheep Blood agar plates and incubated at 37°C under anaerobic conditions (Anoxomat, Advanced Instruments). Culture bottles that remained negative after 14 days were also subcultured under anaerobic conditions and did not result in bacterial growth. All subsequent culturing after initial anaerobic conditions were done aerobically. Two CSF samples were positive for growth in culture bottles. As previously described, colonies from subculture plates were used for Gram stain, organism identification by MALDI-TOF, biochemical testing, and antimicrobial susceptibility testing [13]. Biochemical testing was performed using API 50 CH strip following manufacturers protocol. Susceptibility testing and interpretations were performed by E-test method using Clinical and Laboratory Standards Institute (CLSI) guidelines.

### Bacterial genome sequence and assembly

Bacterial cultures were inoculated in either Luria-Bertani (LB) broth from mono-isolates or anaerobic culture bottles. DNA was extracted using either the Prep Cell Culture DNA Isolation protocol (Bionano Genomics, San Diego, CA) or Zymobiomics DNA kit following the manufacturer’s protocol. DNA samples were then sheared to about 400 bp fragments using the E220 Focused ultrasonicator (Covaris, Woburn, MA), and libraries were prepared with Hyper prep kit (Kapa Biosystems, Wilmington, MA) following the manufacturer’s protocol and sequenced on a MiSeq v3 600 cycle at 10 pM and 15% phiX with 10 million reads per sample. For long-read sequencing, libraries were prepared with 1.5 µg DNA following the Native barcoding genomic DNA (with Exp-NBD104, EXP-NBD-114, and SQK-LSK109) with minor modifications using Kapa or NEB T4 ligase and end-repair enzymes without the use of the FFPE DNA Repair Mix.

Genomes of strains Mbale and NRRL B-4156 were assembled as previously described [22]. The genomes of two additional clinical isolates, Mbale2 and Mbale3, were assembled using the same protocol. Briefly, long reads generated on a MinIon (Oxford Nanopore) were preprocessed using Albacore, assembled with Canu [23] and corrected with Pilon [24].

### Genome annotation and protein comparison

We used the Prokaryotic Genome Annotation Pipeline (PGAP) [25, 26] through RefSeq and RASTtk annotation summarized by the PATRIC database to annotate the three clinical isolates and B-4156 strains. Genome annotation and a functional-based comparison were performed using RASTtk [27] with the default parameters for the *P. thiaminolyticus* Mbale and B-4156 strains. Furthermore, the PATRIC database [28] was used for protein comparisons of the annotations (70% minimal coverage and 30% minimal identity) and the proteomics (50% minimal coverage and 30% minimal identity). Mapping of proteins to the genomic assembly was done using CGView [29]. OrthoVenn2 [30] with default settings was used to compare the predicted proteins with proteins from previously sequenced related bacteria, and Gene Ontology terms for the mapped genes were plotted with ggplot2 [31] in R Statistical Computing Program version 4.0.4. An average nucleotide identity calculator was used to compare the overall nucleotide identity [32]. PilFind using default settings was used to identify the motifs in hypothetical proteins for a transmembrane domain [33]. PHASTER was used to identify specific phage insertions in each genome [34].

### RNA isolation, sequencing and analysis

Bacteria were grown in M9 mineral medium with 0.4% glucose or LB media and samples were prepared at five stages of growth from both cultures to evaluate the span of gene expression. RNA was isolated using the Direct-zol RNA Miniprep kit (Zymo, USA) following the manufacturer’s bead beating and DNase I treatment protocol. RNA was prepared for sequencing with the NEBNext rRNA depletion kit (E7850, New England Biolabs, USA), followed by the Stranded Total RNA Prep (Illumina, USA) following the manufacturer’s protocol. Counts were generated with HTSeq aligned to the respective PGAP annotated genome. Comparisons were done with the Mbale and NRRL B-4156 aligned to the Mbale annotation. Statistical analysis was done with DESeq2 [35] in R.

### Proteomic preparations and analysis of isolates

Three liquid cultures from three separate colonies each of strains Mbale and B-4156 were analyzed concurrently. All samples were digested with trypsin after protein denaturation, reduction, and alkylation essentially as described previously [36, 37]. The labeled peptides were analyzed by nano-scale liquid chromatography coupled with tandem mass spectrometry (nano-LC-MS/MS). The samples were resuspended in solution A (1% formic acid) and 2.5 µl loaded onto a 75 µm i.d. × 50 cm Acclaim^®^ PepMap 100 C18 RSLC column (Thermo-Fisher Scientific) on an EASY nanoLC (Thermo Fisher Scientific) at a constant pressure of 700 bar in solution A. Prior to sample loading the column was equilibrated to solution A for a total of 11 µl at 700 bar pressure. Peptide chromatography was initiated with mobile phase solution A containing 2% solution B (100% acetonitrile, 1% formic acid) for 5 min, then increased to 20% solution B over 100 min, to 32% solution B over 20 min, to 95% solution B over 1 min and held at 95% solution B for 29 min, with a flow rate of 300 nl/min. Data was acquired in data-dependent acquisition (DDA) mode. The full-scan mass spectra were acquired with the Orbitrap mass analyzer with a scan range of m/z = 325 to 1500 and a mass resolving power set to 70,000. Ten data-dependent high-energy collisional dissociations were performed with a mass resolving power set to 17,500, a fixed lower value of m/z 100, an isolation width of 2 Da, and a normalized collision energy setting of 27. The maximum injection time was 60 ms for parent-ion analysis and product-ion analysis. The target ions that were selected for MS/MS were dynamically excluded for 20 sec. The automatic gain control (AGC) was set at a target value of 10^6^ ions for full MS scans and 10^5^ ions for MS2. Peptide ions with charge states of one or > 8 were excluded from MS/MS interrogation.

Data from the mass spectrometer were converted to peak lists and the MS2 spectra were analyzed using Peaks software [38]. The software was used to search a database of each strain’s PGAP annotations with trypsin specificity. A maximum of 3 missed cleavages was allowed. The searches were performed with a fragment ion mass tolerance of 50 ppm and a parent ion tolerance of 25 ppm. Carbamidomethylation of cysteine was specified in Peaks as a fixed modification. Deamidation of asparagine, formation of pyro-glutamic acid from N-terminal glutamine, acetylation of protein N-terminus, oxidation of methionine, and pyro-carbamidomethylation of N-terminal cysteine were specified as variable modifications. Peptide and protein identifications were exported and results were filtered post processing using peptide −10logP values.

### Genome editing with CRISPR-Cas9

All plasmids for genome editing were constructed in E. coli DH5α (fhuA2 lac(del)U169 phoA glnV44 Φ80’ lacZ(del)M15 gyrA96 recA1 relA1 endA1 thi-1 hsdR17) and, after sequencing confirmation, were transformed into E. coli BW29427 (RP4-2(TetS, kan1360::FRT) thrB1004 ΔlacZ58(M15) ΔdapA1341::[erm pir^+^] rpsL(strR, thi-hsdS-pro-) (The Coli Genetic Stock Center) selecting on LB medium containing 50 μg/ml chloramphenicol and 100 ug/ml diaminopimelic acid (DAP). All plasmid constructions were performed with the appropriate enzymes from New England Biolabs. Our genome editing vector was derived from plasmid pJOE9734 (Bacillus Genetic Stock Center) [39], which contains the oriT/traJ origin of transfer, kanR antibiotic resistance gene and cas9 under control of the mannose promoter (PmanPA) from Bacillus subtilis 168 [40]. Furthermore, it carries the pE194ts origin of replication, making its propagation temperature sensitive. A chloramphenicol resistance gene (cat) and its promoter from pAD43-25 were cloned into pJOE9734 at the AfeI site to create plasmid pAS1.

For genome editing to delete the type IV pilus operon, about 400bp DNA upstream and downstream of the T4P operon were amplified separately and ligated by overlap PCR to produce the template sequence for gene deletion by homologous recombination. This DNA product was cloned into pAS1 at *Sfi*I site to generate plasmid pAS2. A guide RNA targeted to Type IV pilus operon was designed with the Integrated DNA Technologies (Coralville, IA) online guide RNAs design tool (https://www.idtdna.com/site/order/designtool/index/CRISPR_PREDESIGN) and cloned into pAS2 at *Bsa*I site to produce pAS3. Plasmid pAS3 was transformed into E. coli BW29427 and a transformant mated with *P. thiaminolyticus* Mbale at 30°C on LB minus DAP overnight and then resuspended in 1 ml LB. Varying volumes were plated on LB plates with 25μg/ml chloramphenicol at 30°C for two days and single colonies were picked for confirmation of successful conjugation by PCR. For cas9 induction, the pAS3-positive colonies were cultured in 10 ml LB with 5% mannose and chloramphenicol at 30°C at 50 rpm for two days, then spread on LB plus chloramphenicol plate and cultured in 39°C incubators overnight to induce loss of pAS3. Resultant colonies were confirmed for deletion of the T4P operon by PCR. Primers used are summarized in **Table S1**.

### Virulence testing using C57BL/6J mice

All animal experiments were performed with oversight by the Penn State Institutional Animal Care and Use Committee, and with Institutional Biosafety Committee approval at biosafety level 2 (BSL2). The virulence of Mbale and Δ(*pilT pilC pilB*) (KO) strains was tested using the mouse infection model as previously described [13]. Briefly, cultures were grown in liquid LB overnight at 37°C and then subcultured onto blood agar plates overnight. Bacteria were harvested from plates and resuspended in saline to a concentration of 2×10^9^ cells/ml. C57BL/6J mice were inoculated with 100 μl of bacterial suspension intraperitoneally and followed for 30 days or until death.

### PCR of *pilT* gene

DNA was isolated as above from either cerebrospinal fluid (CSF) or blood from patients with hydrocephalus. PCR of the *pilT* gene was performed with OneTaq (NEB, M0207) with 500 nM of forward (gatcataatcaatgagcccggtcatgg) and reverse (cttgtgcgaaggcgctgcga) primers in 25 μl reactions. Amplified products were separated on a 1% agarose gel and visualized after staining with ethidium bromide, scoring for the presence of a 200 bp product from *pilT*.

## Results

### Complete assemblies of clinical isolates and type strain B-4156

We previously described the isolation of a novel bacterial strain, which we designated *Paenibacillus thiaminolyticus* Mbale, from the cerebral spinal fluid (CSF) of a patient with post infectious hydrocephalus (PIH) [13] and identified it as *P. thiaminolyticus* based on MALDI-TOF analysis, rRNA similarity, average nucleotide identity and phylogenetic analysis [22]. Two additional isolates, which we designated Mbale2 and Mbale3, were recovered from the CSF of two additional patients with PIH and also identified as *P. thiaminolyticus* by MALDI-TOF analysis. Computerized tomography scans of the brains of the infants from which these strains were recovered are shown in Fig 1A. We previously provided the complete genome assemblies of the Mbale strain and the *P. thiaminolyticus* type strain NRRL B-4156^T^ (B-4156) [22]. The B-4156 strain assembly yielded a complete 6,613-kilobasepair (kbp) chromosome from long-read sequencing, while the initial clinical Mbale strain required optical mapping to construct one complete chromosome from three contigs assembled by the long-read sequencing; three shorter contigs remained unmapped (Fig 1B, Table 1). PHASTER analysis identified 2 of the 3 unmapped contigs as either complete or incomplete phage sequences (**Fig S2B**) [34]. The third unmapped contig likely constitutes an insertion in the chromosome identified by optical genome mapping but flanked by extended repeated sequences, rendering it unmappable by short and long read sequencing. The complete sequence of the two new isolates revealed a single 6,460 kbp contig for the Mbale2 strain and two contigs, 6,561 kbp and 12 kbp, for the Mbale3 strain (Table 1, Fig 1B). PHASTER analysis identified the 12 kbp contig of Mbale3 as an incomplete phage genome (**Fig S2,** Table 1). We further confirmed the species assignment with average nucleotide identity (ANI) [32]. The average two-way ANI of the Mbale, Mbale2, and Mbale3 to B-4156 were 97.05%, 97.03%, and 97.01%, respectively, which fall above 95% sequence similarity across species [41]. Biochemical testing using API test strips read at 48 hours also identified B-4156, Mbale, Mbale2, and Mbale3 as *P. thiaminolyticus* at 99.9%, 99.9%, 97-98.6%, and 91.6-98.9% confidence (**Table S2**).

**Figure 1.**
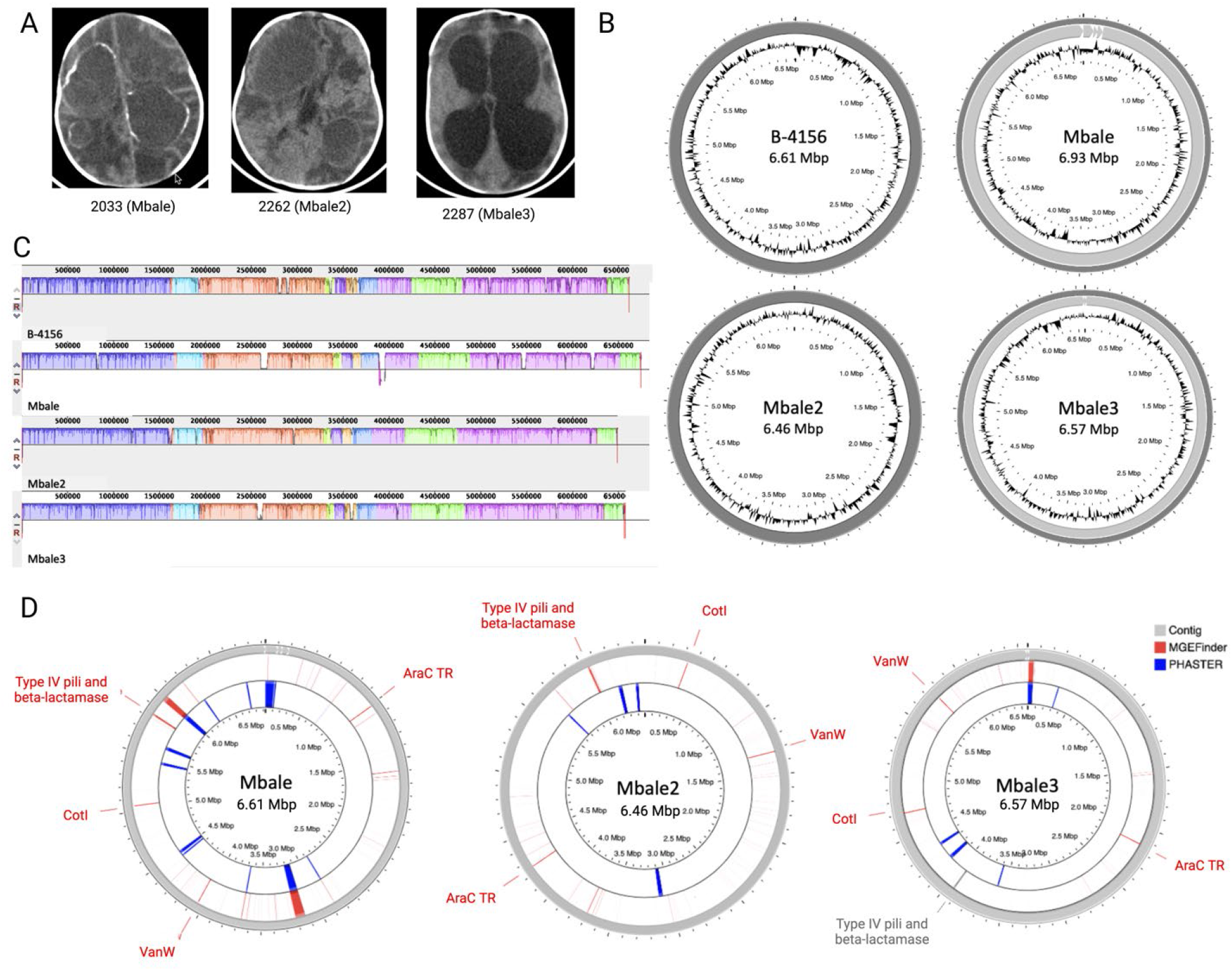
Genomic Characterization of Clinical Isolates of *P. thiaminolyticus*. A) Computerized tomography scans from the three infants with postinfectious hydrocephalus from whose cerebrospinal fluid (CSF) the three clinical isolates were recovered. The first two images from 2033 (clinical isolate strain Mbale) and 2262 (clinical isolate strain Mbale2) were taken prior to surgery and demonstrate loculations and calcified abscess formation. The third image from 2287 (clinical isolate strain Mbale3) was taken after surgery for hydrocephalus and also shows evidence of extensive brain parenchyma damage. B) Shown are complete assemblies of the type strain of *P. thiaminlytocus* B-4156^T^ (B-4156) and three clinical isolates Mbale, Mbale2, and Mbale3, obtained using long and short-read sequencing. The B-4156 and Mbale3 genomes each consists of one continuous contig while the Mbale and Mbale2 genomes each comprises one large contig plus additional contigs (Table 1). C) Alignment with MAUVE of the clinical isolates’ genomes to that of the B-4156 type strain identified twelve locally colinear blocks (LCB), which are indicated by the different colors. White regions within the colored regions represent regions of low sequence similarity. D) Mobile genetic element finder (MGEFinder) identified regions of genomes that could have been derived from mobile genetic elements (MGE, red bars) and could account for the regions of low sequence similarity. Genes specified in red were identified by RAST as being encoded in the predicted MGEs and include the antibiotic-resistant vancomycin gene (VanW), spore coat protein (CotI) and an arabinose family transcription regulator (AraC). Furthermore, a predicted MGE in Mbale and Mbale2 encoded an operon for the type IV pilus, which, although not predicted to be an MGE in Mbale3, was also present in that genome (gray). Regions of phage sequences, identified by PHASTER, are labeled in blue.

**Table 1:**
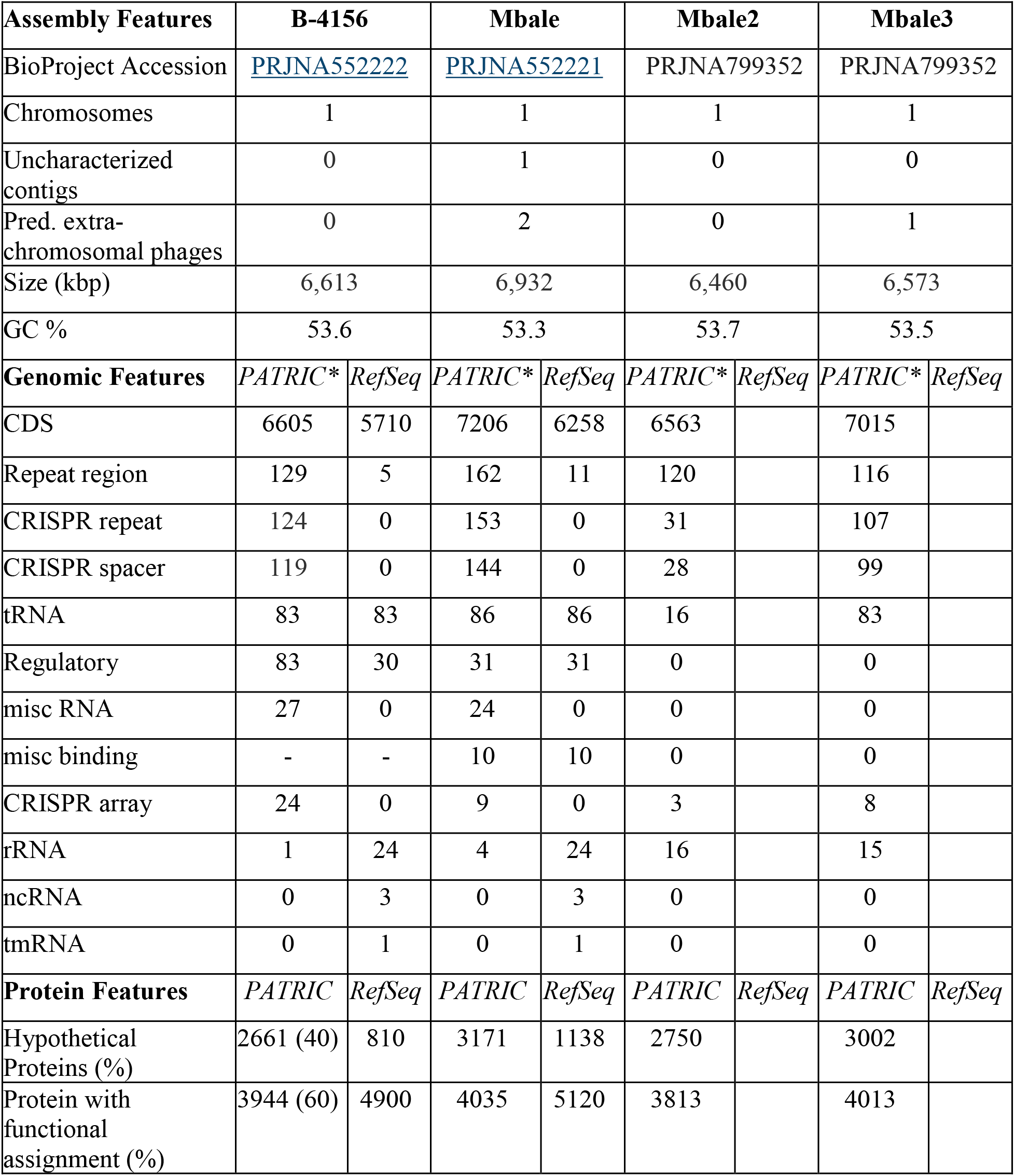
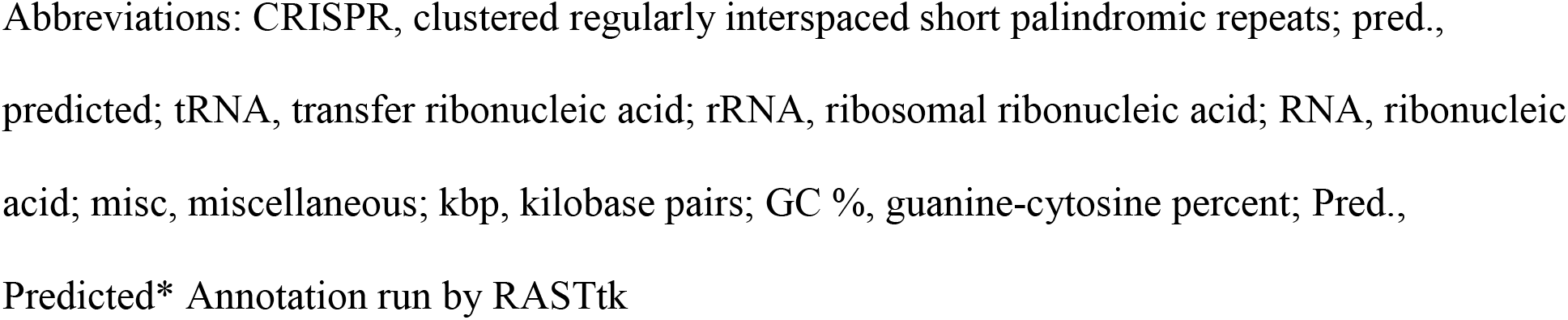
Genomic Feature Summary of *P. thiaminolyticus* Strains. Assembly summary along with Genomic Features of the type strain, B-4156, and clinical strain, Mbale, designated by PARTIC and RefSeq

| Assembly Features | B-4156 |  | Mbale |  | Mbale2 |  | Mbale3 |  |
| --- | --- | --- | --- | --- | --- | --- | --- | --- |
| BioProject Accession | PRJNA552222 |  | PRJNA552221 |  | PRJNA799352 |  | PRJNA799352 |  |
| Chromosomes | 1 |  | 1 |  | 1 |  | 1 |  |
| Uncharacterized contigs | 0 |  | 1 |  | 0 |  | 0 |  |
| Pred. extra-chromosomal phages | 0 |  | 2 |  | 0 |  | 1 |  |
| Size (kbp) | 6,613 |  | 6,932 |  | 6,460 |  | 6,573 |  |
| GC % | 53.6 |  | 53.3 |  | 53.7 |  | 53.5 |  |
| Genomic Features | *PATRIC\** | *RefSeq* | *PATRIC\** | *RefSeq* | *PATRIC\** | *RefSeq* | *PATRIC\** | *RefSeq* |
| CDS | 6605 | 5710 | 7206 | 6258 | 6563 |  | 7015 |  |
| Repeat region | 129 | 5 | 162 | 11 | 120 |  | 116 |  |
| CRISPR repeat | 124 | 0 | 153 | 0 | 31 |  | 107 |  |
| CRISPR spacer | 119 | 0 | 144 | 0 | 28 |  | 99 |  |
| tRNA | 83 | 83 | 86 | 86 | 16 |  | 83 |  |
| Regulatory | 83 | 30 | 31 | 31 | 0 |  | 0 |  |
| misc RNA | 27 | 0 | 24 | 0 | 0 |  | 0 |  |
| misc binding | - | - | 10 | 10 | 0 |  | 0 |  |
| CRISPR array | 24 | 0 | 9 | 0 | 3 |  | 8 |  |
| rRNA | 1 | 24 | 4 | 24 | 16 |  | 15 |  |
| ncRNA | 0 | 3 | 0 | 3 | 0 |  | 0 |  |
| tmRNA | 0 | 1 | 0 | 1 | 0 |  | 0 |  |
| Protein Features | *PATRIC* | *RefSeq* | *PATRIC* | *RefSeq* | *PATRIC* | *RefSeq* | *PATRIC* | *RefSeq* |
| Hypothetical Proteins (%) | 2661 (40) | 810 | 3171 | 1138 | 2750 |  | 3002 |  |
| Protein with functional assignment (%) | 3944 (60) | 4900 | 4035 | 5120 | 3813 |  | 4013 |  |

Abbreviations: CRISPR, clustered regularly interspaced short palindromic repeats; pred., predicted; tRNA, transfer ribonucleic acid; rRNA, ribosomal ribonucleic acid; RNA, ribonucleic acid; misc, miscellaneous; kbp, kilobase pairs; GC %, guanine-cytosine percent; Pred., Predicted\* Annotation run by RASTtk

### Genomic comparisons of *P. thiaminolyticus* B-4156 and clinical isolates

We performed a Progressive Mauve alignment [42] to identify the genomic differences between the clinical isolates and type strain B-4156. The multiple sequence alignment identified 12 locally colinear blocks (LCB) between the four strains, representing sequences with conserved segments and no rearrangements (Fig 1C). The genomes also contained several regions of low sequence similarity. The contigs that did not assemble into the chromosome had no homology to B-4156. The chromosome of the Mbale strain is approximately 132 kb larger than that of B-4156, whereas Mbale2 and Mbale3 genomes are about 153 kbp and 40 kbp smaller than the genome of B-4156 strain, respectively.

Horizontal transfer of mobile genetic elements (MGEs) facilitates rapid evolution of microbial genomes [43, 44]. To investigate genomic differences driven by MGEs, we used Mobile Genetic Element Finder (MGEFinder) [45] and PHASTER [34]. We identified 320 unique sequences and 186 clusters that are identified using CD-HIT-EST at 90% sequence identity over 85% of their sequence per the MGEFinder methods. The 320 unique sequence insertions range from 70-82,113 bp in length in the clinical isolates (median, 168 bp; interquartile range [IQR], 112-352 bp). Mbale, Mbale2, and Mbale3 carry 158, 156, and 129 inserted sequences, respectively, as well as 11, 4, and 5 phage insertions (Fig 1D).

To assess the possible functional significance of the insertion sequences predicted from the MGEFinder, we assessed their coding potential by string analysis and RAST annotations. This analysis yielded 676 predicted coding sequences associated with 22 Gene Ontology terms (**Fig S3**), dominated by genes associated with DNA processing and the establishment of competence to acquire insertion sequences. All three isolates also contained separate insertions carrying genes encoding vancomycin B-type resistance protein VanW, spore coat protein CotI, and the transcription regulator of AraC family (Fig 1D). Antibiotic resistance testing confirmed that the isolates were resistant to vancomycin (**Table S3**). In addition, strains Mbale and Mbale2 carried a common 12 kbp insert spanning a type IV pilus (T4P) operon linked to a beta-lactamase class C-like penicillin binding protein gene. Although not predicted to be an insertion by MGEFinder, Mbale3 also carries a 14 kbp insertion containing the T4P genes identified by annotation linked to a beta-lactamase class C-like penicillin binding protein gene (Fig 1D). Consistent with the presence of a beta-lactamase gene in the isolates, two of the three isolates were resistant to the beta-lactam antibiotics ampicillin and penicillin.

### Predicted proteomes of the clinical isolates and the type strain

The number and classes of proteins predicted to be encoded by each of the three clinical isolates and the type strain, as determined by RefSeq or PATRIC, are listed in Table 1. Consistent with the high similarity of the genomes of clinical isolates and B-4156, these genomes carry very similar sets of metabolic genes and, accordingly, are predicted to encode similar metabolic pathways. These include, among others, similar systems of amino acid biosynthesis, purine, pyrimidine, and cofactor biosynthesis, carbohydrate and lipid metabolism, amino acid and protein degradation, respiration, and energy metabolism (Table 2). Many of the predicted carbohydrate metabolic pathways were confirmed by biochemical metabolism testing (**Table S2**). The clinical isolates and B-4156 all carry a number of antibiotic resistance genes and contain a thiol activated cytolysin, a potential virulence factor. B-4156 carries an operon encoding the type VII secretion, which the clinical isolates lack.

**Table 2:** Predicted Protein by Subsystem from RASTtk Annotation.

| **Subsystems [Subsystem (genes)]** | **B-4156** | **Mbale** | **Mbale2** | **Mbale3** |
| --- | --- | --- | --- | --- |
| **Metabolism** | 91 (745) | 93 (767) | 92 (742) | 94 (768) |
| Cofactors, Vitamins, Prosthetic Groups | 21 (228) | 22 (246) | 20 (231) | 21 (227) |
| Amino Acids and Derivatives | 29 (227) | 29 (225) | 29 (220) | 29 (241) |
| Fatty Acids, Lipids, and Isoprenoids | 11 (88) | 11 (91) | 11 (88) | 11 (92) |
| Carbohydrates | 9 (64) | 8 (65) | 9 (60) | 9 (69) |
| Nucleosides and nucleotides | 8 (64) | 9 (62) | 8 (62) | 4 (27) |
| Iron acquisition and metabolism | 4 (25) | 4 (25) | 4 (28) | 4 (17) |
| Phosphate metabolism | 1 (20) | 2 (21) | 2 (12) | 2 (15) |
| Metabolite damage and mitigation | 4 (17) | 4 (17) | 4 (25) | 2 (7) |
| Secondary metabolism | 2 (7) | 2 (10) | 2 (10) | 2 (5) |
| Nitrogen Metabolism | 1 (4) | 1 (4) | 2 (5) | 2 (5) |
| Sulfur Metabolism | 1 (1) | 1 (1) | 1 (1) | 1 (1) |
| **Protein Processing** | 44 (249) | 43 (255) | 42 (237) | 41 (225) |
| Protein Synthesis | 31 (201) | 31 (203) | 30 (188) | 29 (177) |
| Protein Fate | 13 (48) | 12 (52) | 12 (49) | 12 (48) |
| **Energy** | 28 (205) | 29 (218) | 27 (209) | 29 (235) |
| Energy and Precursor Metabolites Generation | 18 (124) | 20 (141) | 18 (134) | 20 (159) |
| Respiration | 10 (81) | 9 (77) | 9 (75) | 9 (76) |
| **Cellular Processes** | 27 (174) | 27 (183) | 27 (175) | 26 (175) |
| Cell Cycle, Cell Division and Death | 11 (82) | 11 (82) | 13 (82) | 11 (83) |
| Prokaryotic cell differentiation | 13 (77) | 13 (83) | 11 (78) | 12 (77) |
| Mobility and Chemotaxis | 1 (9) | 1 (9) | 1 (9) | 1 (9) |
| Clustering-based subsystems | 1 (5) | 1 (8) | 1 (5) | 1 (5) |
| Microbial communities | 1 (1) | 1 (1) | 1 (1) | 1 (1) |
| **Stress Response, Defense, Virulence** | 36 (120) | 17 (119) | 32 (121) | 31 (124) |
| Resistance to antibiotics and toxic compounds | 19 (66) | 17 (61) | 15 (65) | 15 (124) |
| Stress response: Heat/Cold | 2 (17) | 2 (22) | 2 (18) | 2 (17) |
| Stress response: Osmotic stress | 4 (13) | 7 (13) | 4 (13) | 7 (15) |
| Stress response: Electrophile toxicity | 1 (4) | 1 (4) | 1 (4) | 1 (4) |
| Host-pathogen interactions | 11 (72) | 1 (1) | 1 (1) | 2 (6) |
| **DNA Processing** | 18 (89) | 19 (117) | 18 (90) | 19 (116) |
| DNA Repair | 12 (59) | 13 (96) | 12 (60) | 13 (84) |
| DNA uptake, competence | 2 (16) | 2 (28) | 2 (15) | 2 (16) |
| DNA recombination | 2 (5) | 1 (4) | 2 (6) | 2 (5) |
| DNA replication | 1 (4) | 1 (2) | 1 (4) | 1 (5) |
| **Membrane transport** | 11 (72) | 11 (68) | 7 (63) | 7 (68) |
| ABC Transporters | 1 (42) | 1 (41) | 1 (42) | 1 (41) |
| Cation transporters | 4 (22) | 3 (17) | 3 (17) | 3 (21) |
| Multidrug efflux systems | 2 (3) | 2 (3) | 2 (3) | 2 (4) |
| Protein secretion systems, Type VII | 3 (4) | 0 (0) | 0 (0) | 0 (0) |
| Protein secretion systems, Type II | 1 (1) | 1 (2) | 1 (1) | 1 (2) |
| Uni, Sym, Anti transporter | 0 (0) | 1 (1) | 0 (0) | 0 (0) |
| **RNA Processing** | 12 (52) | 12 (54) | 12 (51) | 2 (16) |
| **Cell Envelope** | 3 (15) | 3 (15) | 3 (14) | 1 (6) |
| **Regulation and cell signaling** | 3 (12) | 3 (12) | 3 (14) | 3 (14) |

As a means of identifying potentially clinically relevant features of the pathogenic strains, we used OrthoVenn2 to compare the predicted proteome across the clinical isolates and B-4156, which identified 6608 unique coding regions across all four strains with 5109 coding regions present in all isolates (Fig 2A). Overall, the clinical strains share more coding regions among themselves than with the type strain (Fig 2B). Specifically, the three clinical isolates share 342 coding regions that are absent from B-4156. Gene ontology of these 342 coding regions returned 39 terms with the largest number of genes associated with “sequence-specific DNA binding”, “plasma membrane”, and “sporulation resulting in the formation of a cellular spore” (**Table S4**). Several genes populate terms such as secretion, iron, and response to toxins that could contribute to the difference in virulence between the isolates and the reference strain (Fig 2C). In addition to these strict paralogous coding regions, each of the three clinical isolates carry a number of unique coding regions, eight of which encode functionally identical or related proteins. These eight regions, versions of which are present in all three clinical isolates but not B-4156, include three genes that encode the T4P, including the twitching motility protein (*pilT*), type IV fimbrial assembly protein (*pilC*), and leader prepilin peptidase (*pilD*) (Table 3). Mbale2 and Mbale3 carry an additional N-methyltransferase related to the T4P. In addition, the clinical isolates carry three phosphorous metabolism enzymes provisionally assigned to the phosphoenolpyruvate phosphomutase subsystem that are responsible for the first step of the carbon-nitrogen bond formation for the synthesis of a broad class of antibiotics (Table 3). The strains also carry single predicted proteins involved in tryptophan and teichoic acid processing. These unique predicted proteins listed in Table 3 were all confirmed in the Mbale strain by proteomic analysis (see below). In sum, our genomic annotation reveals several candidate genes potentially important for pathogenesis, including antibiotic resistance genes and the T4P system.

**Figure 2.**
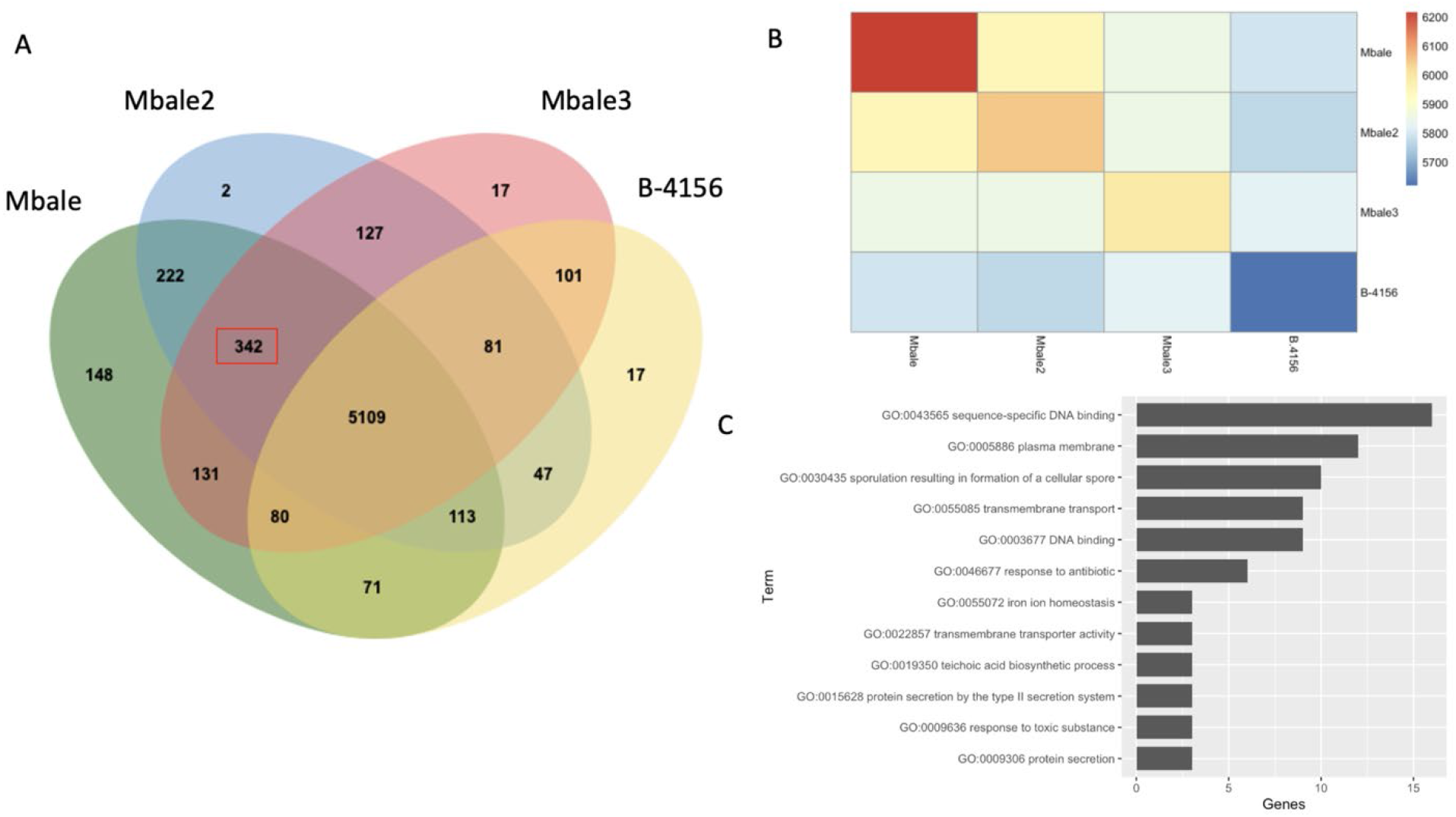
Comparisons of the Predicted Proteomes of the Clinical Isolates and the type strain. A) Venn diagram from OrthoVenn2 analysis comparing the four annotations of the B-4156, Mbale, Mbale2, and Mbale3 strains. The 342 paralogous clusters among the clinical isolates that were absent in the nonpathogenic B-4156 are outlined in red. B) A heatmap quantifying the predicted paralogous clusters across each isolate. C) Clinically relevant gene ontology terms from the 342 proteins that were unique to the clinical isolates.

**Table 3:** Predicted functional proteins in the clinical isolates but absent in the type strain B-4156.

| **Category** | **Predicted Function** |
| --- | --- |
| **Secondary Metabolism** | Phosphoribosylanthranilate isomerase (EC 5.3.1.24) |
| **Phosphorus Metabolism** | Phosphoenolpyruvate phosphomutase (EC 5.4.2.9) |
|  | Phosphoenopyruvate decarboxylase (EC 4.1.1.82) |
|  | 2-aminoethylphosphonate: pyruvate aminotransferase (EC 2.6.1.37) |
| **Membrane Transport & Secretion Systems** | Leader peptidase (Prepilin peptidase) (EC 3.4.23.43) |
|  | Type IV fimbrial assembly protein PilC* |
|  | Twitching motility protein PilT |
| **Cell Wall & Capsule** | Teichoic acid translocation permease protein TagG* |

\*Confirmed with RNASeq and Proteomics

### RNA and Protein Expression Analysis of Mbale and the Type Strains

As an initial step in assessing whether the predicted coding regions are functional, we performed sequence analysis of RNA isolated from Mbale and B-4156 grown in two different media, LB and minimal salts (M9) plus glucose, at five stages of growth (lag phase, middle log phase, late log phase, stationary phase and death phase) (Fig 3A). We identified 6285 and 5817 unique transcripts from the Mbale and B-4156 strains, respectively, with 5893 transcripts present in both strains, 400 unique to Mbale and 8 unique to B-4156 (Fig 3C). Unsupervised hierarchical clustering of the most variable expressed transcripts provided clear discrimination based on strain and stage of growth, with some distinction related to growth media (Fig 3A).

**Figure 3:**
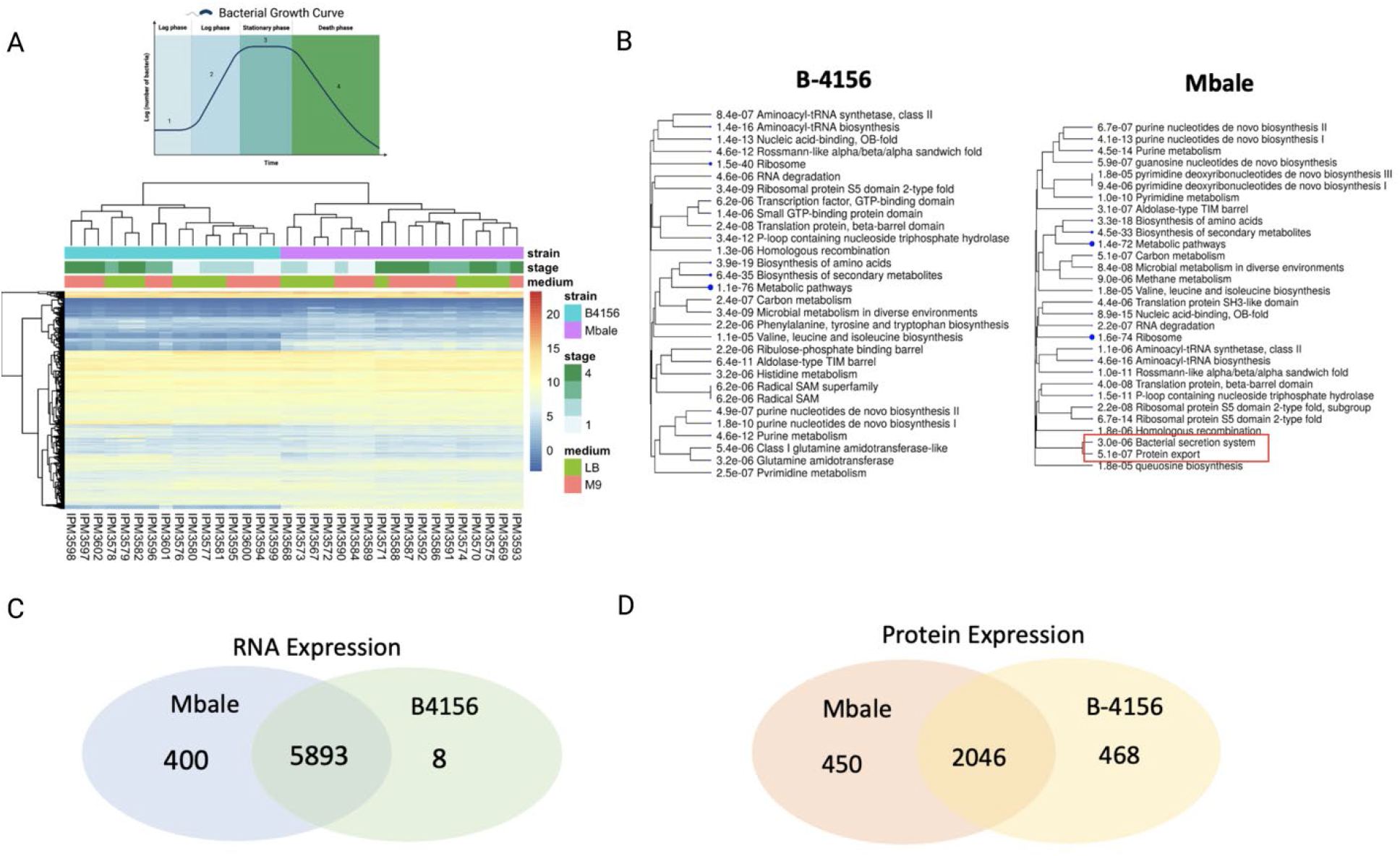
RNA and protein expression of the B-4156 and Mbale strains. A) Culture growth stages (top) and unsupervised hierarchical clustering (bottom) of the RNA transcript levels of the most variable genes expressed in the Mbale and type strains of *P. thiaminolyticus*. Clusters separate primarily according to strains and secondarily on the basis of stage of growth and media. B) Gene ontology of the 450 proteins uniquely identified in proteomic data for the B-4156 strain (left) and the 468 proteins unique to the Mbale strain (right). P-values indicate the extent of enrichment of the genes of each indicated ontological group in the set of genes unique to each of the strains. Bacterial secretion system and protein export genes related to the SecA pathway uniquely represented in the Mbale strain are highlighted in red. C,D) Comparison of RNA (C) and protein (D) expression between the Mbale and B-4156 strains.

To further assess the functionality of predicted proteins, we performed proteomics analysis of the B-4156 and Mbale strains growing exponentially in LB medium. The *de novo* analysis identified 193,878 and 194,234 peptides in the Mbale and B-4156 strains, respectively, which mapped to 2,488 and 2,531 proteins based on PGAP annotation. Two thousand forty-six were present in both strains while 450 unique proteins were expressed in Mbale and 468 in B-4156 (Fig 3D). String analysis of the unique proteins in each strain confirmed the presence of proteins from similar metabolic pathways despite the unique protein sequences. Two unique terms, “Bacterial secretion system” and “Protein export” were identified in the Mbale strain (Fig 3B). The proteins assigned to these terms encode for the SecA secretion pathway (*secY, secA, ffH, laspA, secF, secD*). In addition, the *pilC*-and *tagG*-encoded predicted proteins unique to Mbale were confirmed.

### All three clinical isolates carry a complete T4P operon

The initial annotation of the 12-14 kbp insertion encoding T4P components documented a common gene organization for the element in all three clinical isolates, which encompasses four genes encoding proteins associated with the T4P assembly: *pilD, pilB, pilC and pilT*. This annotation, though, failed to identify genes essential for a complete T4P assembly, namely, those encoding the minor and major pilins and the alignment proteins, PilM, PilO and PilN [46]. However, using BLAST and alignment with COGs [47] and Pfam [48] in comparison to the type II secretion system databases [49–51], we identified a number of these essential assembly genes in the cluster (Fig 4A ,Table 4). Furthermore, we used pilFind to identify three genes that encode for the transmembrane domain on the N-terminus characteristic of pilins (Table 4), one of which had homology to the pseudopilin protein GspH. Comparison of the gene order in the 14 kbp insertion of *P. thiaminolyticus* Mbale with the previously characterized *pil* operons from other Gram-positive bacteria [33, 52] showed the closest match to the *pil* operons of *Clostridium cellulolyticum* H10 and *Bacillus* sp. NRRL B-14911 (**Fig. S4**). The Mbale2 and Mbale3 insertion sequences contained the same gene order present in the Mbale strain (Fig. 3A). In contrast, strain B-4156 encodes only a truncated *pilB* gene (encoding the first 60 amino acids of PilB) and lacks the remaining genes required for complete assembly of the T4P. As shown in Fig 4C, all of these genes in T4P cluster are transcribed in the Mbale strain as is the *pilB* fragment in B-4156.

**Figure 4:**
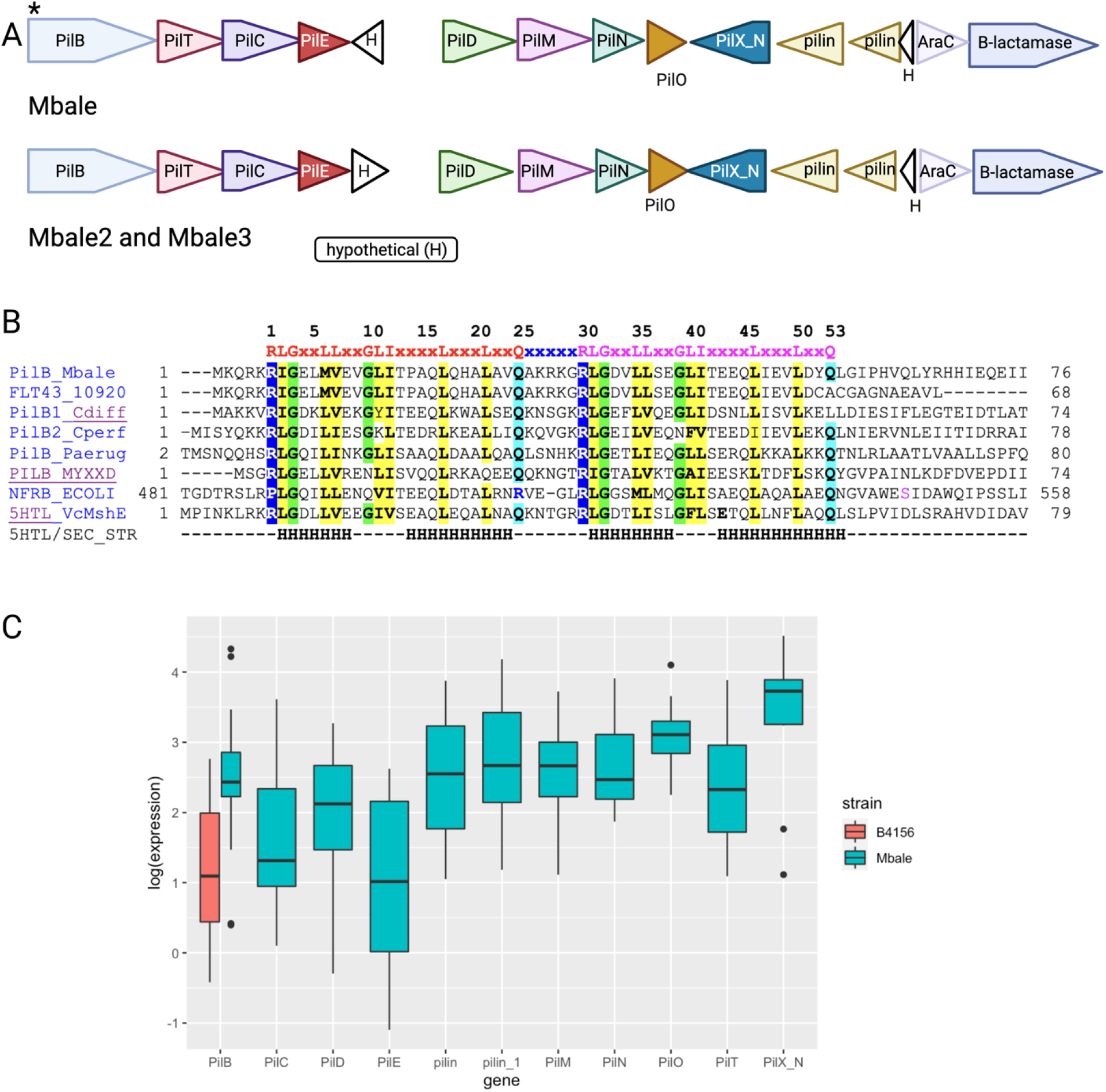
Predicted insertion carries the full Gram-positive T4P operon. A) The T4P operon present in all three clinical isolates and absent in the B-4156 strain located in the predicted mobile genetic element insertion (in strain Mbale, locus tags FLT15_06255 – FLT15_06190). The genes were annotated with RAST or PGAP and/or with hits to the COG and Pfam databases. PilFind [33] was used to identify potential pilins (light brown). White triangles designate hypothetical proteins of unknown function. * indicates a c-di-GMP-binding site in PilB. B) Sequence alignment of the N-terminal fragment of PilB from strain Mbale (GenBank accession NGP58005.1) and truncated protein FLT43_10920 (GenBank: QDM43956.1) from *P. thiaminolyticus* type strain B-4156 against experimentally characterized MshEN domains. The top line shows the conserved c-di-GMP-binding site of the MshEN domain, which consists of tandem 24-aa motifs separated by a 5-aa insert [54]. Aligned sequences include MshEN domains of PilB proteins from *Clostridioides difficile* [82], *Clostridium pefringens* [81] *Pseudomonas aeruginosa* (PA3740) [54, 88] and *Myxococcus xanthus* (MXAN_5788) [89], and from *Escherichia coli* NfrB [90, 91]. The bottom two lines show the sequence and secondary structure (H, α-helix) of the structurally characterized MshEN protein from *Vibrio cholerae* (VC_0405, [54, 92]). Conserved hydrophobic residues are shaded yellow, conserved Gly residues are shaded green. C) Expression levels of the *pil* genes obtained from the RNA-seq data.

**Table 4:** Type IV pili predicted proteins in *Paenibacillus thiaminolyticus* Mbale.

| Protein function | Protein domain (COG, Pfam) | Type II secretion |  | Type IV pili |  |  |  |
| --- | --- | --- | --- | --- | --- | --- | --- |
|  |  | *Escherichia coli* K12 | *Klebsiella oxytoxa* | *Pseudomonas aeruginosa* | *Clostridium perfringens* | *Paenibacillus thiaminolyticus* |  |
|  |  |  |  |  |  | Mbale1 | B-4156 |
| Pilus assembly ATPase | COG2804, PF00437 | GspE | PulE | PilB, PA4526 | PilB, CPE1844, CPE2286 | FLT15_06255, NGP58005.1, 566 aa | FLT43_10920, QDM43956.1, 68 aa |
| Pilus retraction ATPase | COG2805, PF00437 | N/A | N/A | PilT/PilU, PA0395, PA0396 | PilT, CPE1767 | FLT15_06250, NGP58004.1, 362 aa | N/A |
| Core membrane protein | COG1459, PF00482 | GspF | PulF | PilC, PA4527 | PilC, CPE2285, CPE1843 | FLT15_06245, NGP58003.1, 403 aa | N/A |
| Prepilin/pseudopilin | COG2165, PF07963, COG4968 | GspG, GspH, GspI, GspJ, GspK | PulG | PilE/PilV/PilW PA4551, PA4552/56 | PilE, CPE2284 | FLT15_06240, NGP58002.1, 142 aa | N/A |
| Prepilin peptidase | COG1989 | GspO | PulO | PilD, PA4528 | PilD, CPE2287 | FLT15_06230, NGP58000.1, 255 aa | N/A |
| Inner membrane accessory protein | COG4972 | GspL | PulL | PilM, PA5044 | PilM, CPE2283 | FLT15_06225, NGP57999.1, 387 aa | N/A |
| Inner membrane accessory protein | PF05137 | GspM | PulM | PilN, PA5045 | PilN, CPE2282 | FLT15_06220, NGP57998.1, 200 aa | N/A |
| Inner membrane accessory protein | COG3167, PF04350 | N/A | N/A | PilO, PA5046 | PilO, CPE2281, CPE2288 | FLT15_06215, NGP57997.1, 252 aa | N/A |
| Major pilin/pseudopilin | CPOG4969, PF00114 | GspG | PulG | PilA, PA4525 | PilA, CPE2288 | N/A | N/A |
| Outer membrane secretin | COG1450, PF00263 | GspD | PulD | PilQ, PA5048 | N/A | N/A | N/A |
| Pilus Assembly protein | COG4726, PF14341 | N/A | N/A | PilX, PA4553 | PilX_N | FLT15_06210, NGP57996.1, 475 aa | N/A |

The above analysis indicates that the clinical isolates possess the components necessary for assembly of a T4P and that, at least in the Mbale strain, those genes are transcribed as well as translated. This genetic feature of the clinical isolates may account for our anecdotal observation that cells of strain Mbale appear to have higher virulence after growth on agar plates than after growth in liquid culture (data not shown). While we have not rigorously quantified this observation, it is consistent with the reported higher T4P production in the surface-grown *Clostridium perfringens* [53]. In both *C. perfringens* and strain Mbale, the T4P assembly ATPase, PilB, contains an N-terminal c-di-GMP-binding MshEN domain [54] that has been implicated in the regulation of various cellular processes in pathogenesis (Fig. 4B).

### The Mbale T4P system is a virulence factor

To directly test the role of the T4P in virulence, we used CRISPR-Cas9 to construct a variant of the Mbale strain from which the *pilB*, *pilC*, and *pilT* genes were deleted. As described in Materials and Methods, we constructed plasmid pAS3 in which a mannose-inducible promoter controlled Cas9 expression, a sgRNA targeted the region adjacent to the *pilC* gene (Fig 5B) and homology-directed recombination template that would yield a deletion of the three targeted genes (**Fig S5**). Following the introduction and propagation of the plasmid in Mbale, we recovered several colonies from which the three genes were deleted. We assessed the virulence of one of the mutant strains in a mouse infection model (Fig 5A) and determined that, with identical bacterial doses, none of the Mbale injected mice survived while all of the mice injected with the deletion strain survived (Fig 5B, p<0.0001). This result confirms that the T4P system in the Mbale strain contributes to its pathogenesis.

**Figure 5:**
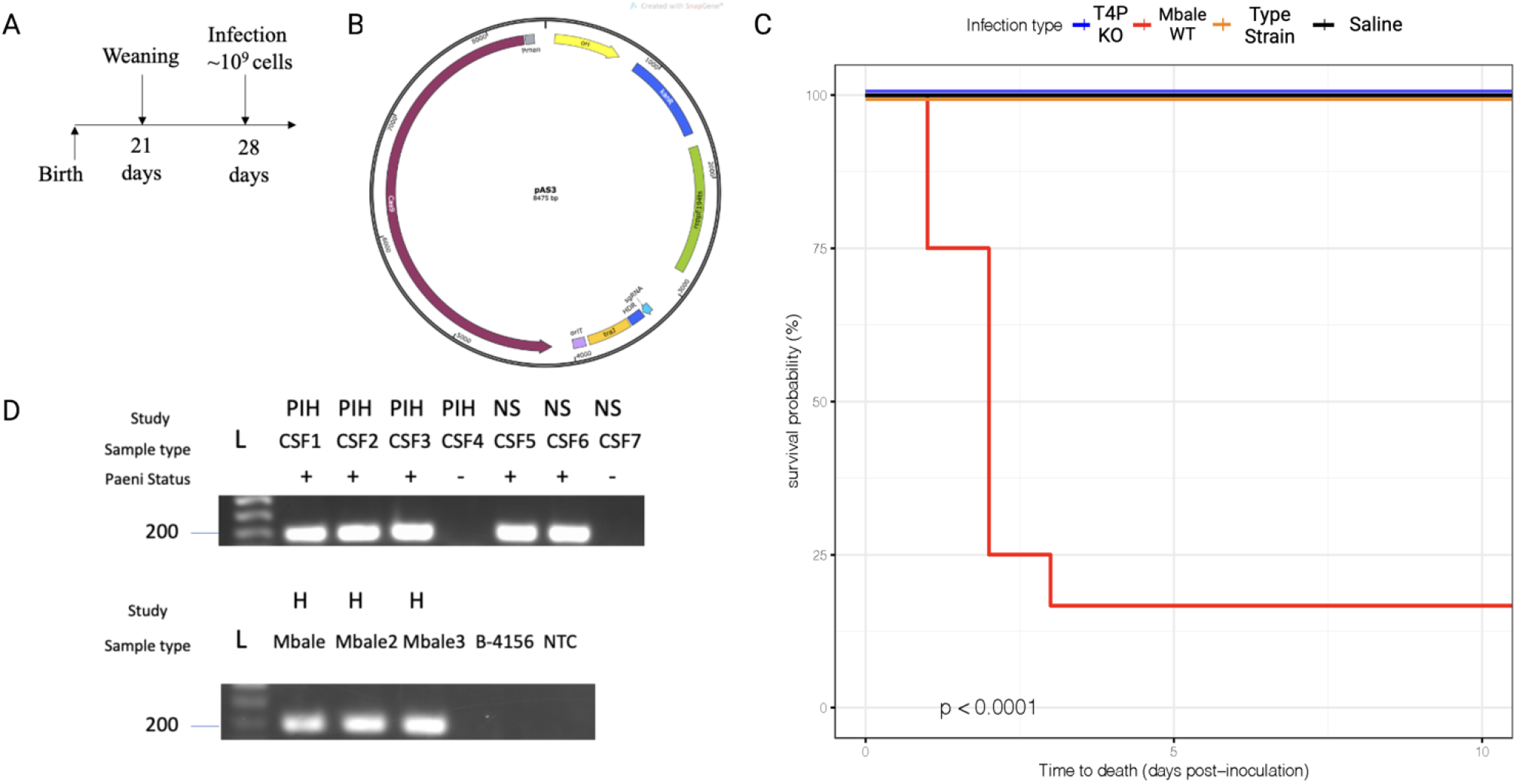
The T4P is a critical virulence factor in all three clinical isolates. A) Mouse sepsis model of infection with injection planned post-weaning on day 28 of life. B) Cas9 containing vector under the control of the mannose promoter used for CRISPR-Cas9 knockout with the homology directed region (HDR), small guide RNA (sgRNA) and kanamycin resistance gene (kanR) for selection. C) Kaplan-Meier survival curve depicting survival of mice following injection of the T4P knockout strain (n=9, blue), the wildtype Mbale strain (n=9, red), the type strain B-4156 strain (n=9, orange) or vehicle control (black) (p<0.0001). D) Amplification of the *pilT* gene in cerebrospinal fluid (CSF) or bacterial culture (H) from infants with postinfectious hydrocephalus (PIH) or neonatal sepsis (NS). *Paenibacillus* status (Paeni status) was determined by qPCR of the thiaminase gene previously defined and was 100% concordant. (Morton). L, DNA ladder.

### *pilT* may serve as a diagnostic marker for *P. thiaminolyticus* infections

We previously identified *P. thiaminolyticus* in the CSF of a subset of patients with PIH or neonatal sepsis (Morton et al, unpublished). To determine whether the T4P is associated with infection, we used PCR to probe for the *pilT* gene, which as noted above is present in all clinical isolates but absent in the reference strain, in clinical samples previously determined to be positive or negative for *Paenibacillus*. As a control, we probed for the *pilT* gene in all three clinical isolates and B-4156 and showed that we could detect it in the clinical strains but not in the type strain (Fig 5D). Previously, we defined *Paenibacillus* infections in patients using qPCR to the thiaminase gene (Morton et al., unpublished). Examining clinical samples, we detected *pilT* in the CSF of all three PIH cases with positive *P. thiaminolyticus* cultures and both neonates with sepsis who subsequently developed PIH with *P. thiaminolyticus* infections. We did not detect *pilT* in CSF samples previously determined to be *Paenibacillus* negative (Fig 5D). These results suggest that the presence of *pilT* correlates with infection by *Paenibacillus*, consistent with a role for the T4P cluster in pathogenesis. Moreover, they suggest that *Paenibacillus* may be detectable in clinical samples prior to the onset of hydrocephalus.

## Discussion

*Paenibacillus* species have been isolated and studied from various sources, particularly in agricultural and industrial settings. The rod-shaped, Gram-positive, endospore-forming aerobic bacteria were initially assigned to the genus *Bacillus* but were subsequently recognized as substantially distinct from other *Bacillus* spp. and assigned to a separate genus *Paenibacillus* (“almost bacillus”) with *Paenibacillus polymyxa* as the type species [55]. Since then, this genus has significantly expanded through assignment of many newly described species and transfer of several other former *Bacillus* species, such as *Bacillus alginolyticus*, *Bacillus glucanolyticus*, and *Bacillus thiaminolyticus* [56]. This genus, along with several others, such as *Brevibacillus* and *Cohnella*, was subsequently assigned to the new family *Paenibacillaceae* within the order Bacillales [57], further separating it from *Bacillus* proper.

We recently identified *P. thiaminolyticus* as a novel pathogen associated with postinfectious hydrocephalus (PIH) and neonatal sepsis (NS) in Ugandan infants. Even though *Paenibacillus* species are highly prevalent in the environment, they had not previously been implicated as a significant human pathogen. Accordingly, we know very little about the mode of pathogenesis of the bacteria or the virulence factors responsible for the severe disease observed in children. However, understanding the virulence of the clinical isolates of *P. thiaminolyticus* provides insight into identification, diagnosis and treatment of infection, information essential for preventing devastating sequelae such as PIH.

Our sequencing, assembly and functional annotation of the genomes of three clinical isolates and the type strain of *P. thiaminolyticus* has provided a comprehensive catalog of the genetic content of these strains and our transcriptomic and proteomic analysis confirmed expression of a significant fraction of the predicted genes (Table 2, Fig 3). While the clinical isolates exhibit more than 97% identity with the type strain (Fig 1B), they differ from the type strain predominantly by the presence of a large number of insertions, likely derived by horizontal transfer of mobile genetic elements (MGEs) (Fig 1C). These insertions are predicted to encode mobile element proteins, hypothetical proteins, multiple AraC family transcriptional regulators, phage proteins and spore coat protein, CotI. The AraC family of transcriptional regulators have been implicated in regulation of proteins of diverse function including virulence factors [58, 59]. All three isolates carry an insertion with the vancomycin B-type resistance gene, *vanW*, and all three isolates are non-susceptible to vancomycin in culture (Fig 1C, **Table S3**). Moreover, all three strains carry a beta-lactamase gene and two of the three isolates exhibit resistance in culture to the beta-lactam antibiotic penicillin and ampicillin. Since the first-line antibiotic regimen recommended by the World Health Organization for neonatal sepsis is ampicillin and gentamicin, these findings indicate that updated guidelines should be considered [60]. Our genomic and sensitivity findings suggest that that ceftriaxone might be an alternative to ampicillin in those infections in which *Paenibacillus* is the suspected causative agent.

Of particular interest with regard to virulence of the clinical isolates, all three strains carry a 12-14 kbp insertion with nearly identical organizations of all the genes necessary for a type IV pilus assembly, in linkage with genes for an AraC transcription regulator and the beta-lactamase c-type penicillin-binding protein (Fig 4A). Our proteomic analysis of the Mbale strain documented expression of genes in this operon (Table 3). Significantly, deletion of several genes within the operon substantially reduced the virulence of the Mbale strain according to pathogenesis test with our mouse model. This strongly suggests that the T4P system contributes to the pathogenesis of the clinical isolates.

T4P are thin bacterial appendages present in various bacteria, including several bacterial pathogens, and have been implicated in an array of functions including cellular adhesion [61, 62], cell mobility [63, 64], protein secretion [65], microcolony/biofilm formation [66–68] and horizontal gene transfer [69]. The T4P were initially observed exclusively in Gram negative bacteria but, with the advent of whole genome sequencing, many conserved components of the T4P have been identified and studied in Gram positive species [33, 70]. The T4P functions through cycles of major (PilA) and some minor pilin protein extension (polymerization) and retraction (depolymerization) driven by the cytoplasmic ATPase assembly pilB and pilT proteins respectively [71]. This cytoplasmic ATPase assembly is usually associated with the inner membrane protein PilC and aligned by the PilM, PilN and PilO proteins [71]. Prepilin (pilin) proteins in the structurally related type II secretion system are delivered to the periplasm via the general secretory (Sec) pathway [72] before being cleaved by the prepilin peptidase, PilD (Fig 3B). Accordingly, it is noteworthy that we observed expression of Sec pathway proteins in the clinical Mbale strain but not in the B-4156 strain. Finally, function of the pilus is often defined by the pilin protein structure, which has been characterized as IVa and IVb subtypes [73, 74]. The IVa T4P has generally been associated with eukaryotic cell adhesion [61, 62] and horizontal gene transfer [75] while the IVb pili promote self-adhesion [76, 77]. Accordingly, further study of the structure of the Mbale pilin protein will facilitate characterization of the function of the T4P of in *P. thiaminolyticus* and its role in pathogenesis.

The T4P in our clinical isolates is closely related to those in *Clostridium cellulolyticum* H10 and *Bacillus* sp. NRRL B-14911 (**Fig S4**). The latter strain was originally isolated from the Gulf of Mexico [78] but was subsequently reclassified as *Bacillus infantis* [79]. Notably, *B. infantis* was isolated from a case of neonatal sepsis in Busan, South Korea [80], an observation that reinforces the possible connection of T4P and neonatal sepsis. Moreover, we noted that the PilB protein of our clinical isolates, but not that of the type strain, contains a c-di-GMP-binding motif domain (Fig 4B). In *C. perfringens*, the formation of T4P is controlled in a c-di-GMP-dependent manner [81], which suggests the same might be true for the clinical isolates of *P. thiaminolyticus.* In *Clostridioides difficile*, increased c-di-GMP production of T4P promoted adherence to epithelial cells [82]. A detailed analysis in the regulation of T4P expression in clinical isolates of *P. thiaminolyticus* is expected to provide further insights into the potential contribution of c-di-GMP to the virulence of the clinical isolates.

T4P is a widespread virulence factor that is structurally conserved, which makes it an attractive diagnostic and therapeutic target. Previously, Barnier et al [83] showed that the T4P mediated twitching motility by the PilT protein in the gram-negative pathogen *Neisseria meningitis* is required for sustained bacteremia. Moreover, they showed that adjunctive treatment with phenothiazines targeted T4P reduced vascular colonization and associated inflammation in a mouse model [84]. The impact of the phenothiazine family of drugs on the gram-positive T4P warrants further investigation and could provide a valuable adjunctive therapeutic approach to treatment of neonatal sepsis and PIH caused by *P. thiaminolyticus*. Successful pathogenic bacteria must propagate and survive within a host, which likely requires more that just one virulence factor. By identifying differences in the clinical strains, we have uncovered several candidate genes that may be involved in pathogenesis (Fig 2A). However, focusing on only differences between the clinical isolates and type strain B-4156 may miss certain VFs, such as toxins, that could be present in both pathogenic and nonpathogenic bacteria but only mobilized in the former due inability of the nonpathogenic bacteria to survive in the host or to deliver the toxin [65, 77]. For instance, our clinical isolates as well as the type strain encode a thiol-dependent cytolysin with 84% similarity to alveolysin, the toxin in *Paenibacillus alvei* that can lyse and inactivate eukaryotic cells [85–87]. Further functional genetic analysis of the clinical isolates will be required to identify additional relevant VFs. For instance, we demonstrated that comparative proteogenomic of differentially virulent bacterium of the same species can identify critical VFs that cannot be identified from annotation alone. Specifically, we pinpointed the T4P genes as critical for virulence in the clinical isolates of *P. thiaminolyticus* associated with PIH and NS and showed that they were acquired via a common MGE. This methodology thus provides an unbiased framework to identify key VFs in the bacterium that were previously unrecognized as a significant pathogen.

## Acknowledgments

We thank the Penn State College of Medicine Genomics Core for Illumina sequencing of the bacteria and Dr. Yuka Imamura for her generosity in sharing equipment and materials for MinIon sequencing. We would also like to thank Dr. David Craft for his expertise and guidance on culturing and clinical evaluation of bacteria from clinical samples. The expert technical assistance of James Malone, Dr. Yiling Mi and Rose Connors is gratefully acknowledged. The proteomic experiments were performed at the Washington University Proteomics Shared Resource (WU-PSR), R Reid Townsend MD.PhD., Director and Robert Sprung, PhD., Co-Director). The WU-PSR is supported in part by the WU Institute of Clinical and Translational Sciences (NCATS UL1 TR000448), the Mass Spectrometry Research Resource (NIGMS P41 GM103422; R24GM136766) and the Siteman Comprehensive Cancer Center Support Grant (NCI P30 CA091842).

## Funding

U.S. National Institutes of Health (N.I.H) Director’s Pioneer Award 5DP1HD086071 and NIH Director’s Transformative Award 1R01AI145057. IT and MYG were supported by the Intramural Research Program of the U.S. National Library of Medicine at the NIH.

## Notes

### Competing Interest Statement

The authors have declared no competing interest.

